# Hyperbolic odorant mixtures as a basis for more efficient signaling between flowering plants and bees

**DOI:** 10.1101/2020.05.13.093864

**Authors:** Majid Ghaninia, Yuansheng Zhou, Anina C. Knauer, Florian P. Schiestl, Tatyana O. Sharpee, Brian H. Smith

## Abstract

Animals use odors in many natural contexts, for example, for finding mates or food, or signaling danger. Most analyses of natural odors search for either the most meaningful components of a natural odor mixture, or they use linear metrics to analyze the mixture compositions. However, we have recently shown that the physical space for complex mixtures is ‘hyperbolic’, meaning that there are certain combination of variables that have a disproportionately large impact on perception and that these variables have specific interpretation in terms of metabolic processes taking place inside the flower and fruit that produce the odors. Here we show that the statistics of odorants and odorant mixtures produced by inflorescences (*Brassica rapa*) are also better described with a hyperbolic rather than a linear metric, and that combinations of odorants in the hyperbolic space are better predictors of the nectar and pollen resources sought by bee pollinators than the standard Euclidian combinations. We also show that honey bee and bumble bee antennae can detect most components of the *B. rapa* odor space, and the strength of responses correlates with positions of odorants in the hyperbolic space. In sum, a hyperbolic representation can be used to guide investigation of how information is represented at different levels of processing in the CNS.

## Introduction

Natural odors are typically mixtures of many different chemical compounds (Raguso, 2008). To better understand how these different odorant compounds might be perceived and processed by insects and mammals, it is very useful to work within a low-dimensional representation of odorants (Keller et al., 2017; Ravia et al., 2020; Snitz et al., 2013), where the dimensions define common physical or perceptual properties. At the same time, there is strong evidence in the olfactory system for nonlinear interactions between odors, which suggests that other types of spaces need to be evaluated. Recently, we showed that volatile molecules present in the natural environment produced by fruits can be organized as an approximately two-dimensional map where distances between odors are determined according to a *hyperbolic* metric (Zhou et al., 2018). Hyperbolic spaces are characterized by an exponential expansion of states and provide an approximation to hierarchical tree-like structures (Krioukov et al., 2010). Positions close to the origin describe more central nodes, whereas those further from the origin describe derivative products of reactions in metabolic pathways.

This hyperbolic structure makes intuitive sense for semiochemicals produced by plants. The reason for this is that different chemical compounds often arise from the same or linked metabolic pathways within the plants (Borghi et al., 2017). And so, positions of volatile molecules reflects, and can be used to infer, the activity of underlying metabolic pathways. Thus, taking into account the low-dimensional and hyperbolic aspects of the space of natural odors makes it possible to extract more reliable messages about the state of the plant important to pollinators compared to the case where mixtures are evaluated according to a Euclidean metric.

We focus here on floral odors produced by *Brassica rapa*, which is pollinated by insects such as honey bees (*Apis mellifera*) and bumble bees (*Bombus* sp) that use floral odors to identify flowers that contain nectar and pollen. Floral perfumes of *B. rapa* contain at least 26 different chemical compounds (Kobayashi et al., 2012). We show that bees can detect many or all of the monomolecular odorants in the mixture, and that several can contribute to detection of the mixture. However, the mixture gives a stronger, more reliable signal across a broader range of doses than the individual components. We show that dimensions of the mixtures computed within hyperbolic space had a greater predictive power about the amount of pollen and nectar than those computed using a Euclidean metric. Our data are consistent with a model in which components of hyperbolic submixtures together amplify important, independent messages to bees about floral properties (Zhou et al., 2018).

## Materials and methods

### Experimental model and subject details

In this study we used foragers of two bee species, honey bees (*Apis mellifera*) and bumble bees (*Bombus terristris*). The honey bee foragers were collected at the entrance of the hives located in the School of Life Sciences at Arizona State University, on Tempe campus, AZ, USA. The bumble bee foragers were captured from the entrance of the commercially purchased boxes (Koppert Biological Systems; koppertus.com).

### Preparation and electroantennographic detection

The collected foragers were immobilized by cooling and individually restrained in a row of eight truncated pipette tips ensuring that the heads and the antennae protruded from the narrow end of the pipettes. One antenna of the restrained bee was then cut off from the pedicel and subsequently transferred to a custom-made antennal holder composed of two capillary glass (tip and reference) electrodes (World Precision Instruments, Inc., USA) filled with insect saline solution. Each electrode was positioned alongside of the length of a microscope slide (75 × 25 × 1 mm) and secured on a small ball of dental wax. The tip and the cut base of the antenna were gently mounted between the tip and the reference electrodes, respectively (Supplementary Fig S1). The holder was then placed on the stage of an Olympus BX51WIF light microscope and viewed at 60× magnification for electroantennogram (EAG) experiments.

The EAG experiments were performed by inserting silver wires into the electrodes until the electrical recording with OSNs was attained. The EAG signal was amplified using an AC/DC differential amplifier (A-M Systems, Inc., Model 3000, USA), subsequently digitized using a digitizer (1440 A DigiData, Molecular Devices, USA), visualized and saved on a PC computer, and the maximum depth of the EAG deflection was measured offline using Clampfit 10.3 software. The EAG responses were normalized (and presented in percent) to the control (hexane) to compensate for the plausible antennal fatigue caused by e.g. repeated delivery of various compounds and/or prolonged recording. For normalization, the EAG responses elicited by individual compounds at a given dose were divided by the average response obtained by the control passed over the antenna at the outset and end of each run.

### Odorants

One objective of our study was to evaluate honey bee and bumble bee antennae for sensitivity to a broader array of odorants identified from *B. rapa*. We selected twelve compounds associated with *B. rapa* scent that were previously identified using chemical analyses (Schiestl, 2014; Taveira et al., 2009). This subset included two odorants – phenylacetaldehyde and α-farnesene - identified as important in behavioral studies (Knauer & Schiestl, 2015) as well as other inflorescence and plant related compounds. This subset is not a comprehensive set of chemicals identified from *B. rapa*. But it is sufficient to test for broad sensitivity to chemicals from different classes emitted by the plant and its inflorescences. Odorants were 1-butene-4-isothiocyanate, α-farnesene, methyl salicylate, indole, methyl benzoate, benzyl nitrile, (*Z*)-3-hexenyl acetate, p-anisaldehyde, phenylacetaldehyde, decanal, nonanal, acetophenone. For the mixture we used just a 1:1 combination of the 12 odorants rather than trying to simulate a more natural *B. rapa* mixture. Again, the purpose here was to evaluate how robust the signal from a mixture, rather than specifically the natural mixture, would be relative to the individual odorant components. Other characteristics of the compounds are displayed in Table 1. All compounds were diluted in hexane (Fisher Scientific, USA).

**TABLE 1:**
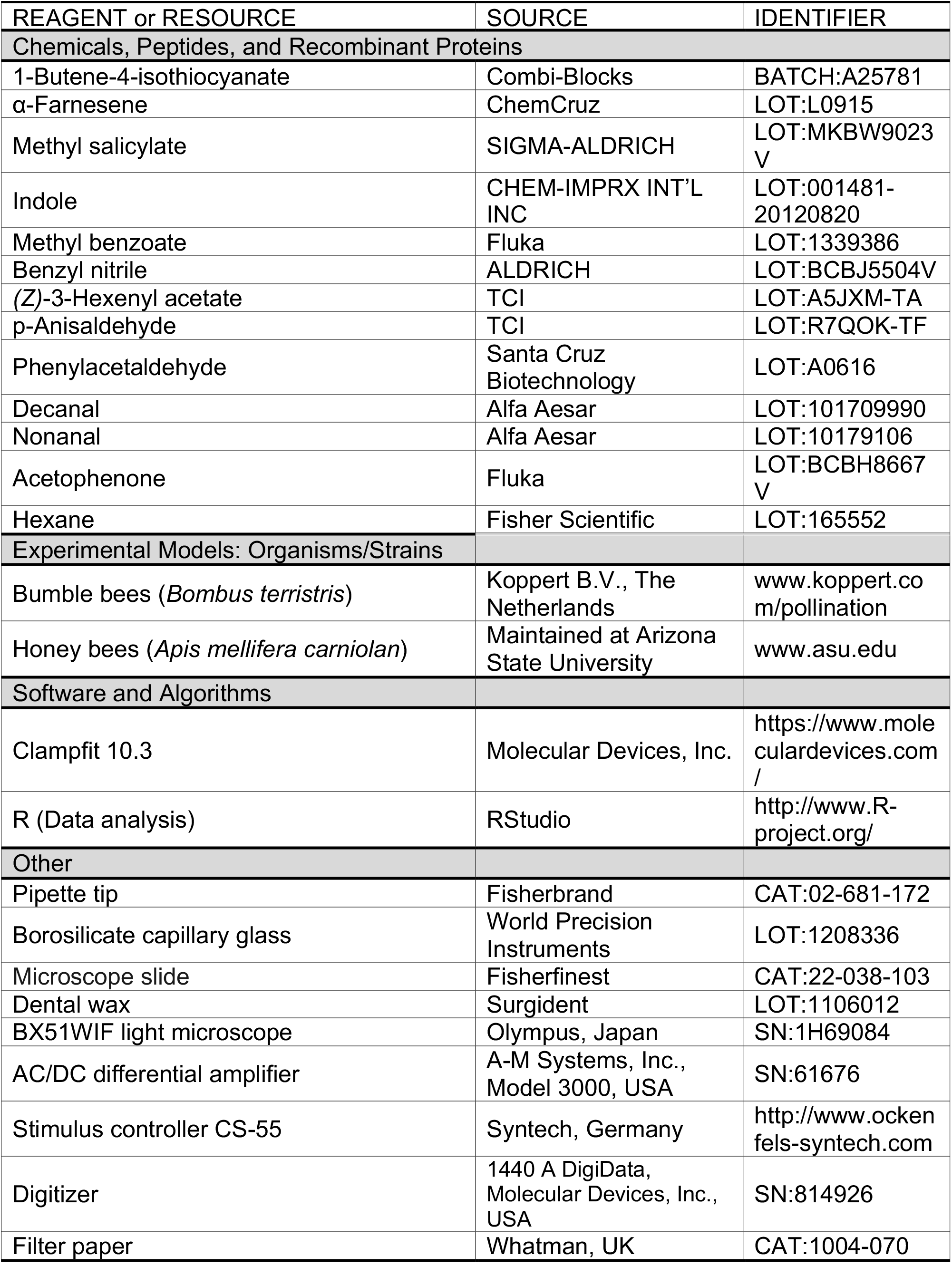
A comprehensive list of all materials and resources used in the manuscript.

### Odor delivery

A 15μl aliquot of a solution of the individual compounds was subsequently loaded onto two pieces of circular filter papers (10 mm diameter each, Whatman, UK) located in the tip of the Pasteur pipettes. The pipette tip was placed into a hole made along a delivery glass tube whose one end was hooked up to a stimulus controller (CS-55, Syntech, The Netherlands) through a Teflon tube and the other end was directed towards the preparation approximately 1-2cm from it. The stimulus controller delivered the individual stimuli for 0.5 sec into a constant 89ml/min humidified and purified airstream passing over the preparation though the delivery glass tube. For every run, a new antennal preparation was used. To prevent the previously delivered odorant molecules from lingering in the antennal periphery, residual odors were evacualted from the arena for 30 sec after each odor delivery, and then another 30sec was allowed to pass (1 minute total time interval) before delivering the next compound. To evaluate whether response declines initially observed at higher doses are due to sensory adaptation persisting over the one minute recovery interval, we repeated the experiment at high doses using a 5 minute interval.

### Dose response experiment

All compounds were applied at increasing doses in linear steps ranging from 10^−11^ g/L to 10^3^ g/L. Different doses of each stimulus were tested only once (always from the lowest to highest doses) on individual antennae. Each dose was tested at least three times and each time a new antennal preparation was used. For the complete mixture, 50μl of each of the twelve compounds at a given dose were combined in a vial so that, at the end, the vial contained a total volume of 600μl (50μl of twelve compounds) at that dose. Therefore, the composite concentration of the mixture is the same as in the individual solutions, but the concentration of individual compounds in that mixture is only 1/12th of that in the individual solutions. Delivery of the compounds and analysis of the responses were done as described above.

### Omission (subtraction) experiment

For the omission experiment, we removed individual compounds from the 10g/L complete mixture. The removal of a given compound in the mixture was compensated for by adding an additional 4.5μl of each of the 11 remaining compounds. In other words, to reach the total volume of 600μl, 54.5μl of each of the 11 remaining compounds (i.e. complete mixture minus compound X) were combined in a separate vial. Therefore, the composite concentration was the same as in the individual solutions, but the concentration of any single compound was only 1/11th of what it is in the individual solutions. In total 13 recordings from both species were made and for each recording a new antennal preparation/individual was used. After each recording, stimuli were randomized. Delivery of the compounds and analysis of the responses were done as described above.

### Floral odor sampling and analytical procedures

Odor samples from B. rapa inflorescences used herein were collected and first published in Knauer & Schiestl (Knauer & Schiestl, 2015), where a description of methods can be found.

### Software

Clampfit 10.3 was used to analyze the intensity of the EAG responses in mV. The software may be available online at www.moleculardevices.com. We used R 3.3.0 for data entry and statistical analysis.

### Statistics

For the dose-response experiments we used data collected at 10g/L as a reference point and applied both the Kruskal-Wallis test and ANOVA on both honey bees and bumble bees to compare the EAG response intensities for each of the *Brassica* odors. We used Tukey HSD post hoc test to confirm where the difference occurred between odorants. For the subtraction experiments we first applied both the Kruskal-Wallis test and ANOVA to compare the responses elicited by the incomplete mixtures (mixture with individual compounds omitted). P-values lower than 0.05 were considered to be statistically significant. All results are represented as mean ± SEM. The ‘‘n’’ represents the number of antennae tested.

### Multi-dimensional Scaling

There are two general categories of performing multi-dimensional scaling: a metric multi-dimensional scaling and a non-metric multi-dimensional scaling (Kruskal, 1964). Metric MDS attempts to approximate the geometric distances *dij* within a low dimensional represention using the loss function defined as:

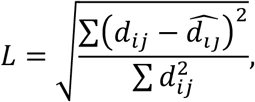

where 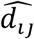 are distances in the low-dimensional embedding. The non-metric MDS by comparison only reconstructs original distances up to a monotonic transformation (only rank-ordering is preserved) (Kruskal, 1964). This confers the algorithm with extra robustness. In this work we used a non-metric MDS while embedding points into a low-dimensional hyperbolic space and using a hyperbolic metric to evaluate distances between data points within the low-dimensional embedding space.

There are multiple representations of a hyperbolic space. Here we used the so-called native representation with polar coordinates (Krioukov et al., 2010). The angular coordinates in this space are the same as in an Euclidean polar coordinates, while the radius R characterizes the hierarchical depth of the structure and measures the degree of hierarchy in data. The distance between two points is evaluated as:

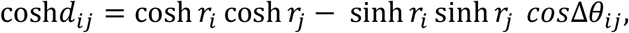

Where ri and rj are the radial coordinates of the two points, and Δ*θ_ij_* is the angle between them. In D-dimensional HMDS, we initialize the embedding process by uniformly sampling points within radius R. The directions of points are uniformly sampled around the high-dimensional sphere, and the radial coordinate is sampled according to the following probability distribution:

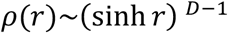

During the iteration process, we update both angular and radial coordinates according to the gradient descent.

## Results

We first evaluated whether honey bees and bumble bees can detect a broad array of chemical compounds of *B. rapa* floral scent. The measurements were made based on electroantennogram (EAG) responses to each of the compounds and to a complete synthetic mixture (Ignell & Hansson, 2004; Olsson & Hansson, 2013). The EAG response to any odorant represents summed excitatory and inhibitory activity across all of the axons from thousands of Olfactory Sensory Neurons spaced along a bee’s antenna (Supl Fig 1). Although EAG measurements are less sensitive than single sensillum recordings, they provide a more appropriate measure for determining whether OSNs on an antenna are collectively capable of detecting a large number of chemical compounds, and EAG’s provide a means to assess relative sensitivity to different odorants. The use of single sensillum recordings would be prohibitive for establishing a complete representation of chemical compounds an antenna can detect because of the large number of recordings that would be needed.

### Honey bees and bumble bees can detect most chemical compounds of the *B. rapa* mixture, which also gives the most robust response

We tested each antenna from honey bees and bumble bees with an increasing dose series of odorants diluted in hexane (Figs 1A and B; Supl Fig 2). We observed no tangible response at doses up to 10^−1^ g/L, at which point responses to some odorants began to increase. By 10^−1^ or 10 g/L both species’ antennae responded to all odorants, with the exception of 1-butene-4-isothiocyanate in honey bees. At higher doses, i.e. 10^2^ and 10^3^ g/L, responses to most pure odorants declined. This decline was likely due to sensory adaptation that persisted for at least the minute that lapsed between each stimulation. The sole exception was phenylacetaldehyde in honey bees, which showed no evidence of adaptation through 10^3^ g/L. We performed further EAG experiments on honey bees to determine how the timing between odor deliveries may have affected the responses to the highest (10^2^ and 10^3^ g/L) doses. We verified that when the OSNs are allowed enough time, i.e. five minutes, their responsiveness failed to show adaptation (Fig 1A).

**Fig 1.**
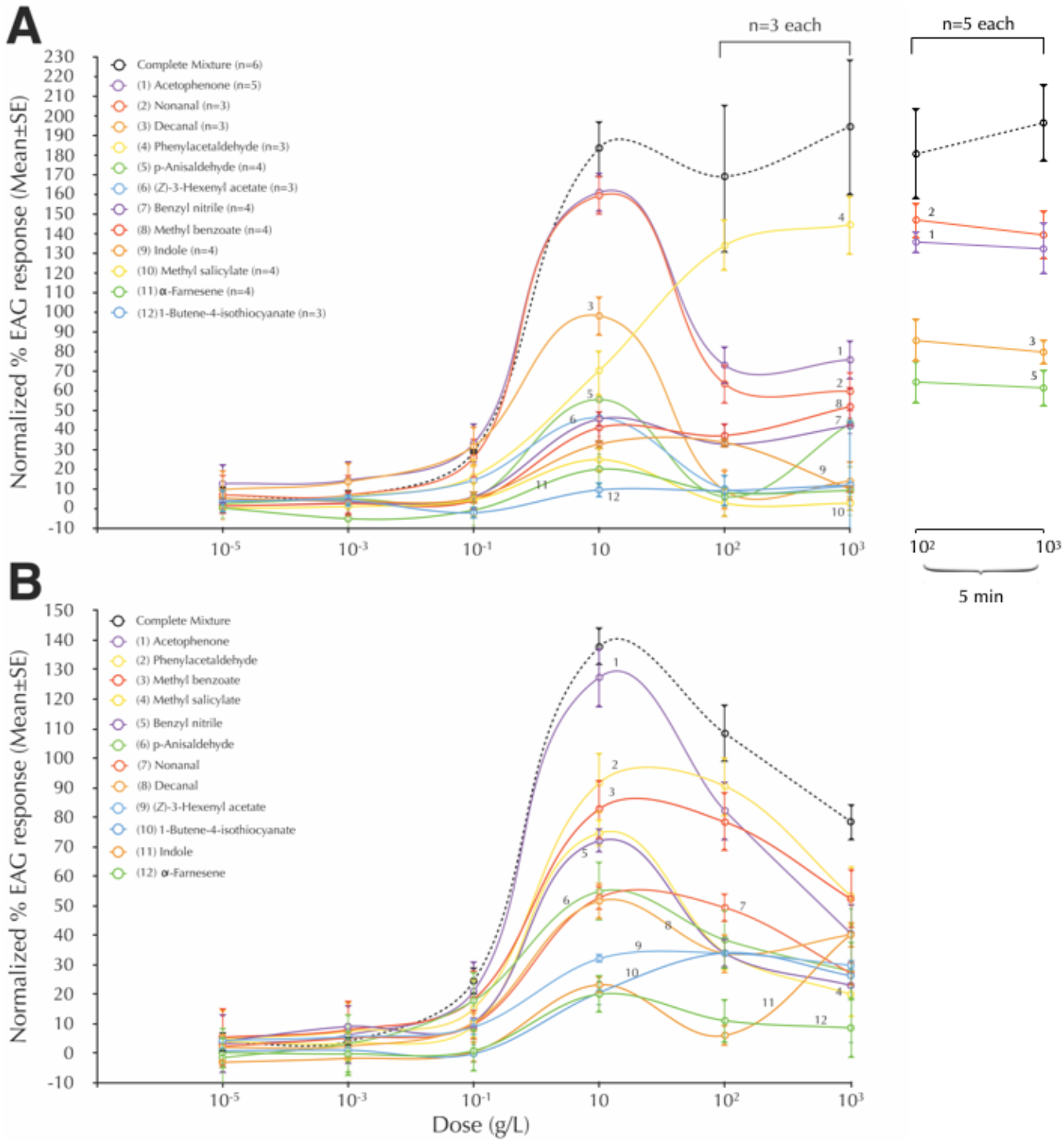
Honey bees and bumble bees can detect almost all components of the *Brassica* mixture, but the mixture gives the most robust response. For the mixture, the x-axis represents total dose of solutes. Electroantennographic responses of (A) honey bees (mean±SE) and (B) bumble bees (mean±SE, n=3) to the selected odorant components of *Brassica rapa* scent. Note that the olfactory responses to most stimuli at 10^2^ and 10^3^ g/L are lower than those at 10 g/L. The responses do not show such a decline after longer inter-test intervals, suggesting that decline with the shorter test interval was due to adaptation (right line-graphs in A). See Supl Fig 2 for statistical analysis.

It is worth noting that in both species the mixture elicited the most robust responses from 10 through 10^3^ g/L. The dose for the mixture refers to the sum of all 12 components. Thus the dose of individual components in the mixture was only a fraction, approximately one-twelfth, of what was presented for each pure odorant at that dose. There was little evidence of adaptation in honey bees to mixtures, even at the short interstimulus intervals. Adaptation was evident in bumble bees; nevertheless, even in this case, the mixture always elicited the highest response.

### Most components contribute to detection of the mixture

To parse out the significance of specific components to the mixture we omitted individual components from the mixture at 10 g/L (see Method Details). The logic was that the EAG response would decrease if a component contributes in any way – either qualitatively or quantitatively via, for example, intensity - to sensory processing of the mixture. Otherwise the response would be unchanged when the component is deleted.

We found that several components contribute to detection of the mixture (Fig. 2). In both species, removal of one component reduced the median normalized response to the mixture. The extent and degree of reduction differed across compounds. In honey bees (Fig 2A), removal of phenylacetaldehyde, nonanal, decanal, acetophenone, and p-anisaldehyde all significantly reduced responses to the mixture. In bumble bees (Fig 2B), removal of all single components except 1-butene-4-isothiocyanate and nonanal significantly reduced responsiveness. However, even in cases where the response did not reach significance, median values and in a few cases the 75% range, were below the normalized mixture response. Thus, it is possible that even in those cases there is some contribution to the mixture, and it would be prudent to include those odors in a broader statistical analysis.

**Fig 2.**
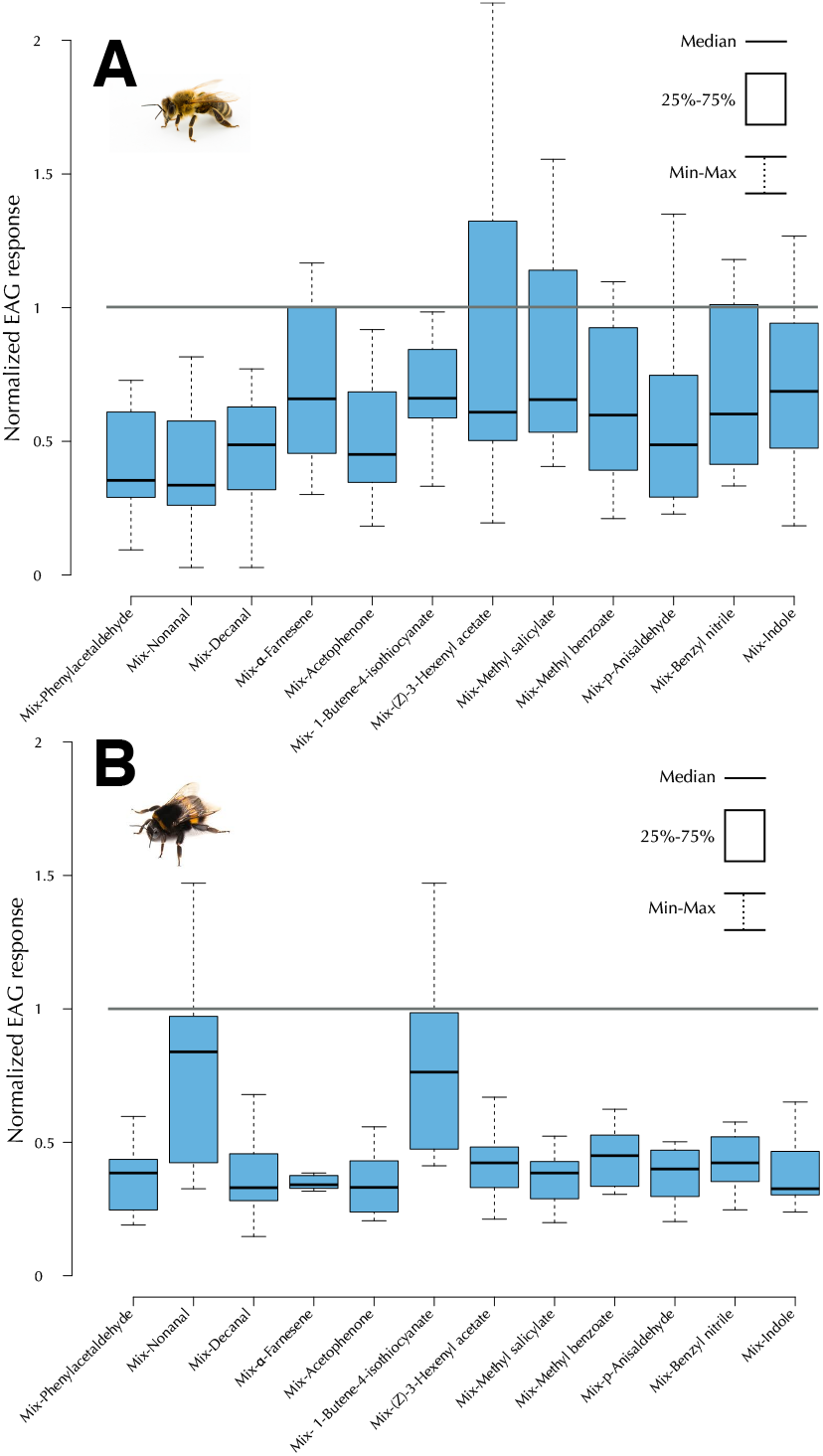
Several components contribute to detection of the mixture. EAG responses of honey bee (mean±SE, n=11) and bumble bee antennae (mean±SE, n=6) to the mixtures of *Brassica rapa* scent with individual component compounds omitted. (A) In *Apis mellifera* omission of phenylacetaldehyde, nonanal, decanal, acetophenone, and p-anisaldehyde from the blend significantly decreased the responses elicited by the OSNs. (B) In *Bombus terrestris* ommission of all compounds, except 1-butene-4-isothiocyanate and nonanal, from the blend significantly reduced the responsiveness of the OSNs compared to the complete mixture. Significant reductions were based on 95% Confidence Intervals not overlapping with 1.0, which was the normalized response to the mixture.

### Submixtures represent nectar and pollen content

Since honey bees and bumble bees can perceive most if not all odorants in the *B. rapa* mixture, we sought to establish the statistical relationships among the odorants and how they correlate to important variables for bees, such as nectar and pollen. Different approaches have been used to perform this kind of comparison. Previous work by Knauer and Schiestl (Knauer & Schiestl, 2015) identified two volatile compounds, phenylacetaldehyde and α-farnesene, that are positively associated with nectar and pollen. We used the same data set to investigate the possibility that submixtures may provide more complete information about nectar and pollen in *B. rapa* florets.

To find additional odor components that are associated with the nectar and pollen content of the flower, and which may supplement detection of those important resources for the bees, we performed two types of analyses. The first analysis was performed with standard Principal Component Analysis using the logarithms of odorant doses. This analysis identified combinations of odorants – i.e. submixtures - that account for largest variance across samples (Fig 3). The top three PC’s cumulatively explained 94.3 of the total odor variance across the *B. rapa* florets and via factor loadings (p≤0.01) collectively accounted for all of the individual odorants. PC1 explained 76.3.% of the variance and was positively associated with benzyl nitrile (0.96) and p-anisaldehyde (0.25). This PC was positively associated with log(pollen)-to-log(nectar) ratio. The second PC explained 17% of the variance and could be associated with the same two odorants but in different proportions and signs: p-anisaldehyde (0.94) and benzyl nitrile (−0.26). This PC component was not correlated, in a statistically significant way, with any linear combination of log(pollen) or log(nectar) content. [We will see that correlations will improve once we transition to the hyperbolic space]. The third PC accounted for 3% of the variance and was positively associated with log(nectar with p=0.02). These components had the highest loading factor: Z-3-Hexenyl acetate (0.67), followed by methyl salicylate (0.55), 1-Butene-4-isothiocyanate (0.29), α-farnesene (0.17), methyl benzoate (0.14), phenlyacetaldehyde (0.13), acetophenone (0.11) and nonanal (0.11).

**Fig 3.**
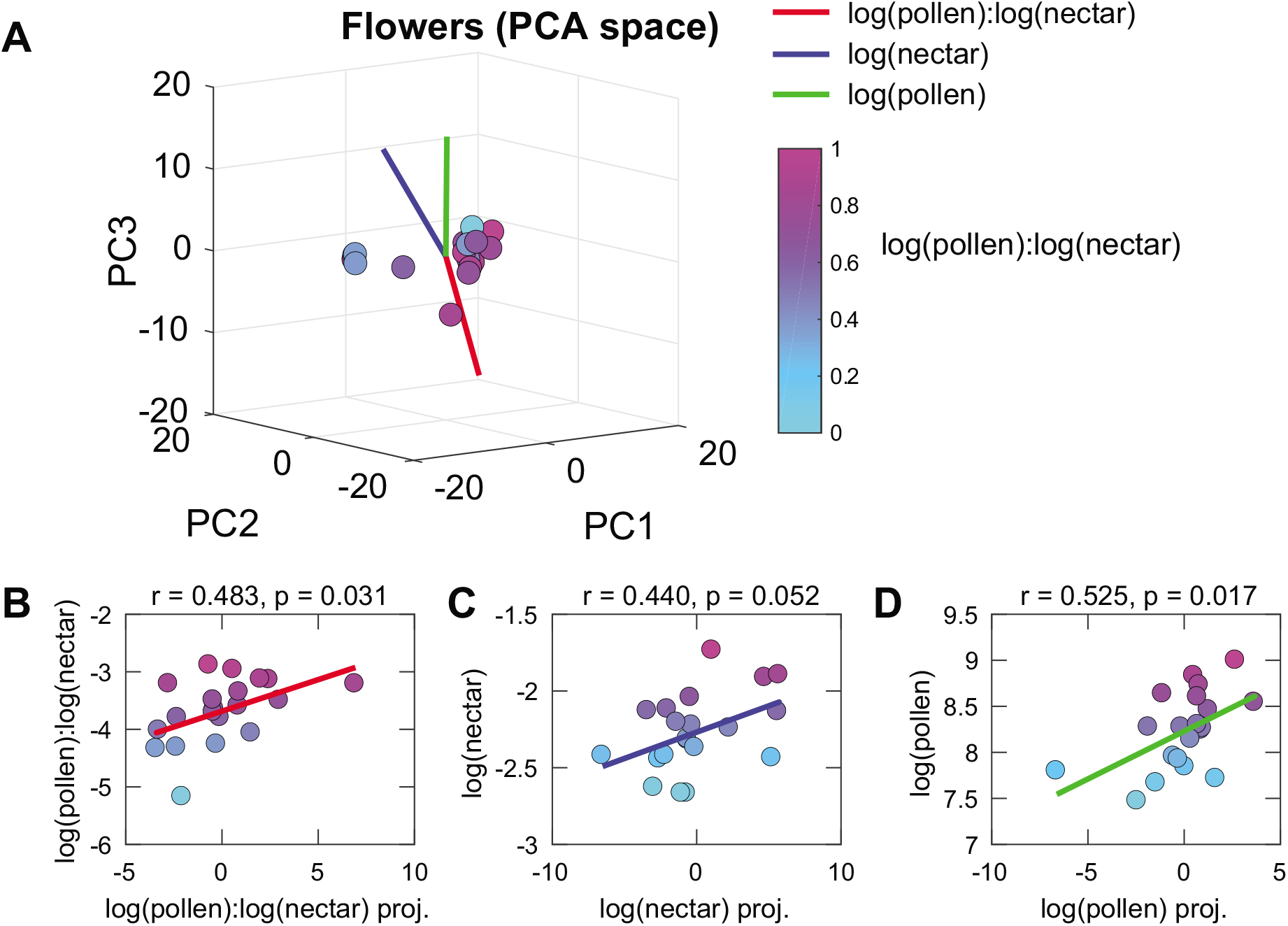
Mixtures of monomolecular odorants that predict pollen and nectar concetrations in the PCA space. (**A**) Representation of odor mixutres produced by Brassica flowers within the space of three leading principal componds (PCs). Axes that are most correlated with florescence nectar and pollen content are shown in red for log(pollen)-to-log(nectar), blue for log(nectar), and green for log(pollen). (B) Correlation with log(pollen)-to-log(nectar) ratio; (C) with log(nectar), and (D) with log(pollen).

### A hyperbolic space as a model of odor predictive power

The PCA analysis assumes a Euclidian metric of the odor space (Koulakov et al., 2011), meaning that changes in the concentration of individual monomolecular odorants have the same impact on the predicted distances between the resulting mixtures regardless of the initial concentration and/or odor identity. We recently published a report that the odor spaces for different types of plant and animal derived volatiles are better fitted using a curved ‘hyperbolic’ metric (Zhou et al., 2018). With this metric the relevant impact of a change in the concentration of an odorant component depends on its initial concentration in ways that cannot be fully captured simply by working with logarithms of the odorant concentration. The impact of odorant concentration change also depended on its identity. The intuition for the relevance of the hyperbolic metric for natural odorants is that the hyperbolic metric approximates activity of hierarchical tree-like processes (Krioukov et al., 2010). Metabolic networks that produce natural odor mixtures certainly have such characteristics. We therefore used hyperbolic Multi-Dimensional Scaling to embed points from *B. rapa* floral volatiles onto a 3D hyperbolic space. The curvature of the spaces was the same as in (Zhou et al., 2018) with R_max_ = 7 and R_min_ = 0.9.

Embedding odorants into this hyperbolic space produced strong correlations with nectar and pollen values (Fig 4). In Fig 4A, circles represent individual monomolecular odorants, and stars represent mixtures produced by individual inflorescences. Distances between these points in hyperbolic space reflect the correlations between odorants or mixtures across samples. Points that are more closely clustered tend to be more correlated across samples. As a first step, we compared the sensitivity of EAG responses to various monomolecular odorants with the odorant positions in the hyperbolic space (Fig 4B). Here, we observed that monomolecular odorants that evoked the strongest EAG responses in honey bees (acetophenone #1, nonanal #2, and decanal #3 cf. Figure 1) have the largest component along the axis that best discriminates between pollen and nectar content (Fig 4A). The rest of the odorants were located in the opposite hemi-sphere and covered it approximately uniformly. Overall, there was a strong correlation between the position of odorants in the hyperbolic space and the EAG responses in antennae. Thus, honey bees show enhanced sensitivity to odorants that best distinguish between pollen and nectar content of inflorescences. The correlation of EAG amplitudes to nectar/pollen in bumble bees was less clear than for honey bees. Except, like in honey bees, the antennae were most sensitive to acetophenone, which is correlated to this reward axis.

**Fig 4.**
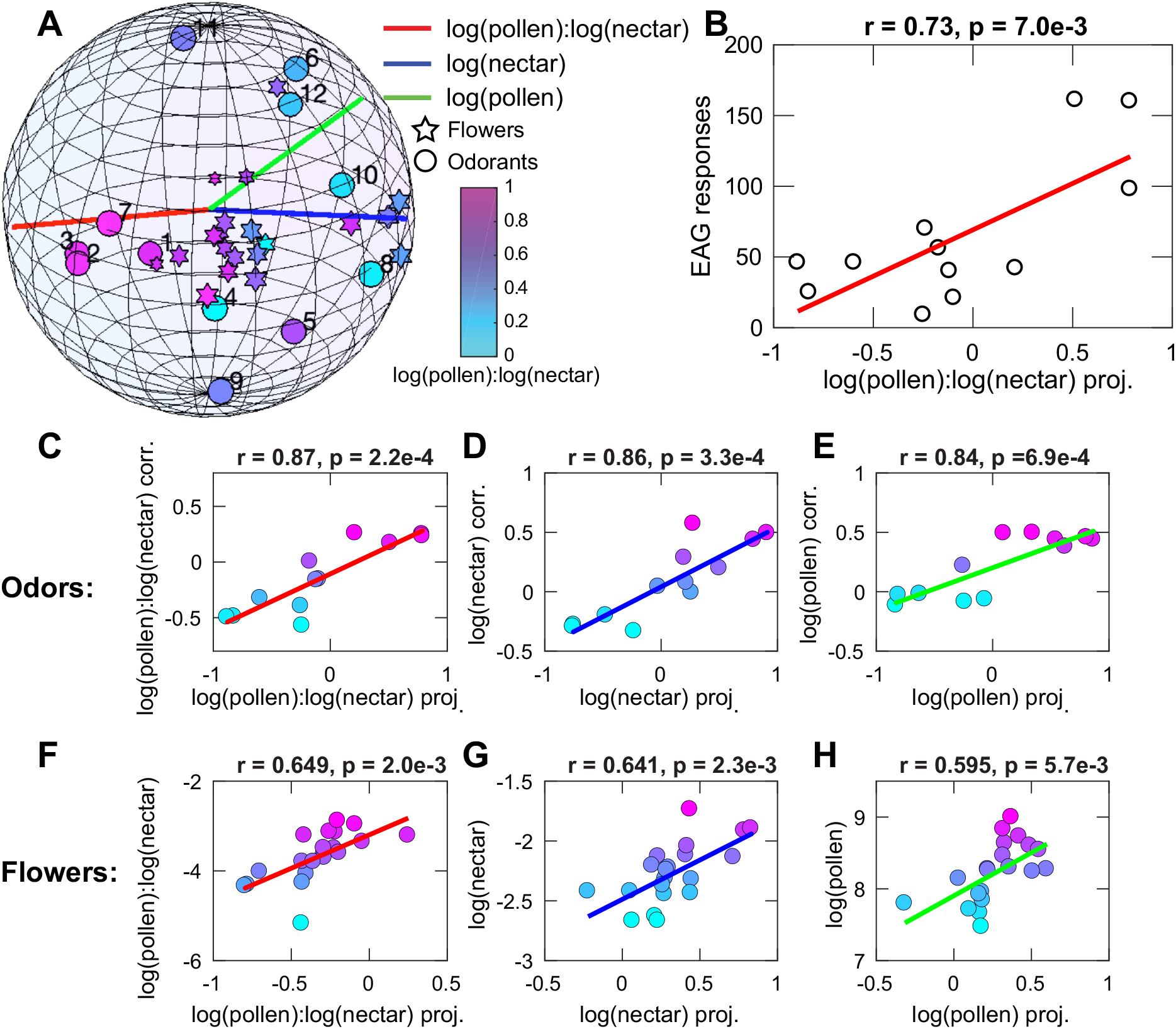
Hyperbolic representation of monomolecular odorants and flower odor mixtures. **(A)** Poincare ball representation of the hyperbolic space (Zhou et al., 2018) together with axes more strongly correlated with log-nectar (blue), log-pollen (green), and their ratio (red). Circles represent individual monomolecular odorants, and stars represent odor mixtures produced by individual inflorescences. Numbers denote odorant identity according to Figure 1A (honey bees). Color denotes the corresponding ratio between log-pollen and log-nectar. For inflorescences this is based on direct measurement, and for odorants on the correlation with their concentration. See also Suppl Figure 3 for a three dimensional version of this figure. (B) Correlation between the strength of EAG response in honey bees (from Figure 1) and odorant position in the hyperbolic space. (C-H) The predicted correlations between projections onto these axes and nectar/pollen measurements.

Odorants and mixtures associated with inflorescences had predictive power for the pollen and nectar content. In particular, we could identify three axes that were strongly correlated with the log-nectar (red), log-pollen (blue) and the ratio between log-nectar and log-pollen (green). The corresponding correlations between projections onto these axes and nectar and pollen flower content are shown for components odorants in Fig. 4C-E and for mixtures from inflorescences in Figs. 4F-H. Here again one can identify three axes that are associated respectively with the log-nectar, log-pollen, and their ratio doses (Figs 4F-H). The correlations are all stronger and more significant than those obtained in the Euclidean space using the PCA analysis (Figure 5). These results indicate that hyperbolic mixtures of monomolecular odorants are better predictors of pollen/nectar flower content than any Euclidean combinations of odorants, which have been identified assuming independent contributions of monomolecular odorants.

## Discussion

Many odorants and odorant mixtures serve as communication signals within and between species. In humans, molecular features of odorants as well as mixtures of odorants are important for determining perceptual qualities that relate to fruit pleasantness and food quality (Gilbert et al., 2015; Schwieterman et al., 2014; Zhou et al., 2018). In fact, there is reason to expect that many fundamental properties of olfaction will apply across a broad phylogenetic spectrum. For example, the honey bee and fruit fly have 163 and 63 functional receptors in their genomes, respectively. (Robertson & Wanner, 2006; Robertson et al., 2003). With the combinatorial nature of receptor responses to any odorant (Hallem et al., 2004), which insects share with mammals (Nara et al., 2011), the perceptual space for odor coding is potentially enormous (Gerkin & Castro, 2015). Therefore, it is a reasonable hypothesis that many olfactory systems have evolved to analyze and parse different meanings out of complex natural odorant mixtures. We propose that understanding the relationships between natural odors and the sensory systems that have evolved to detect them can be aided by a statistical understanding of the structure of the odor landscape, much in the same way that similar analyses have been important for understanding vision (Bell & Sejnowski, 1997; Ruderman & Bialek, 1994) and audition (Lewicki, 2002).

In our previous study (Zhou et al., 2018), we used published data from volatiles across several fruit varieties and mouse urine together with data from human psychophysical ratings (Gilbert et al., 2015; Schwieterman et al., 2014). A hyperbolic space provided better fits for both the volatiles and psychophysical preference data. The hyperbolic nature of the physical odor space has only been explored to date in relation to human detection (Koulakov et al., 2011; Zhou et al., 2018), although indications of hyperbolic structure were present also in earlier results (Ravia et al., 2020; Snitz et al., 2013).

Here we use a beneficial insect-plant system related to pollination to test whether hypotheses that stem from this logic might be more general. *B. rapa* is pollinated naturally by honey bees, bumble bees and hover flies. When pollinated exclusively by bumble bees over several generations, the volatile profile of *B. rapa* is increased quantitatively relative to when pollinated by hover flies, which use more visual than olfactory cues (Gervasi & Schiestl, 2017; Schiestl et al., 2018). This pollinator-based change in odor profile shows the importance of odors in insects that have well-developed olfactory systems, and it implies that there is a significant cost to odor production, which places pressure to make signalling more efficient. Earlier analysis of the same dataset we used here identified a small subset of chemical components, especially phenylacetaldehyde and α-farnesene, that were reliable and – because of the cost - *honest* signals of nectar and/or pollen (Knauer & Schiestl, 2015). Our analyses confirm that both odorants are detectable by honey bee and bumble bee olfactory sensory neurons in the antennae. Moreover, phenylacetaldehyde may also be more resistant to adaptation.

In addition, our analyses show that there is potentially more information present in the relationships between mixture components. The PCA and hyperbolic MDS analyses both indicate that combinations of odorants can serve as indicators of the amount of nectar or pollen in a flower. However, the two methods arrive at that conclusion via different assumptions. PCA assumes independence among the elements – odorants in this case – and it extracts from high to low the linear combinations that explain the most variance in the data set. This method produces correlations with pollen and nectar, but some of the more important odorants identified by Knauer & Schiestl (Knauer & Schiestl, 2015) only load on the third component that accounts for only a fraction of the total variance. While bumble bees learn to pick out whatever component might indicate pollen and/or nectar (Knauer & Schiestl, 2015), it seems as though PCA is not the best method for analysis of floral chemistry because it almost missed two known key components.

In contrast, the hyperbolic space provided a substantially better fit for the nectar and pollen in inflorescences, most likely because of the underlying correlations that arise from the metabolic pathways that produce odorants in the *B. rapa* mixture. Some mixtures and some components were better correlated to pollen, nectar, or both. In our EAG analyses, both the concentration responses and the omission experiment show that several or most of the components of the mixture can be detected by the antennae of both honey bees and bumble bees. We do not at this point know what the biologically relevant range for these compounds might be. However, even if the biologically relevant range is below the detectable limit with EAG or more sensitive measures, these odors could act synergistically to enhance detection of other components of the blend, as we discuss below. Therefore, because the antenna is sensitive to several odorants in the blend, the use of a hyperbolic multidimensional space to analyze the odor profile of *B. rapa* may be important for understanding how this information is represented in the CNS. In fact, sensory processing in both species may be adapted to the information content of *B. rapa* inflorescences. In our data, EAG responses in honey bees were correlated with the position of the odorants in the hyperbolic space, and in both bee species, the strongest EAG responses were for odorants that had strong projections onto the axis in the hyperbolic space that represented the pollen nectar ratio. We also show that phenylacetaldehyde, which is a reliable signal of nectar and pollen in *B. rapa* (Knauer & Schiestl, 2015), elicits strong EAG responses in both species, and it results in less adaptation in honey bees.

Both odorant components and mixtures from inflorescences correlate to nectar and pollen in hyperbolic space. So what is more important, individual components or mixtures? There are several reasons to favor mixtures. In our experiments the mixture of all 12 components, each at a fraction of the dose of the components, elicited the strongest response. Furthermore, at doses above 10 g/l, antennae showed strong sensory adaptation to components. In honey bees the mixture did not elicit adaptation. In bumble bees the mixture elicited adaptation, but responses to the mixture were still the stronger than components at higher doses in spite of adaptation. With a mixture, more different receptor types are targeted, exposing each individual receptor type to lower doses of its cognate chemical compounds. Thus, less adaptation leads to more robust signaling over a larger range of doses. Furthermore, due to non-linear competitive binding of different molecular components to odorant receptors, odor-evoked activity patterns are more stable and first-spike latencies are shorter for mixtures than for pure odorants (Chan et al., 2018). Both properties – less adaptation and enhanced stability - would enhance the accuracy and speed of detectability of mixtures over pure odorants. Competitive binding in mixtures can also predict more complex properties of sensory receptor responses, such as synergy, antagonism and overshadowing (Reddy et al., 2018; Singh et al., 2019). More studies are now needed to investigate how other components may synergize signals from important components like phenylacetaldehyde (Knauer & Schiestl, 2015), for example.

The hyperbolic space, in contrast to PCA, predicts that mixtures can have disproportional impact on perception. Improved predictions from the use of hyperbolic metric that we find here for *B. rapa* inflorescences indicates nonlinear synergistic interactions associated with the presence of multiple odorants. Put differently, in a hyperbolic space the presence of two components will exponentially increase the strength of the cue compared to the presence of individual components. Honey bees and bumble bees searching for pollen, nectar or both could focus on *B. rapa* odor submixtures that independently reflect levels of each reward. Our work also suggests further studies that would, for example, reveal the basis for correlations among components of submixtures – e.g. common biochemical pathways, genetic pleiotropy, epigenetic factors or linkage disequilibrium among genes that produce odor mixture components.

In summary, we have found that hyperbolic representation of odorants reveal stronger association with both flower resources and bee antennal responses, compared to predictions that can be obtained using a Euclidean representation. Hyperbolic representations have several nonlinear aspects that can be particularly useful for characterizing olfaction and natural mixtures of odorants. First, the hyperbolic structure makes it possible to quantify hierarchical relationships in the data. As a result odorants that are correlated with several other odorants, or are more shared across species (Biasazin et al., 2019; Sinding et al., 2013), will be assigned more central positions in the space. Second, based on the properties of vector addition in a hyperbolic space, one expects odorants with more central positions to have a stronger impact within a mixture compared to odorants with more distal positions. From this perspective, it is interesting to note previous reports of stronger tuning in flies for those volatiles that are shared among the plant host species (Biasazin et al., 2019). It will now be fruitful to test the model using other coevolved odor/animal relationships, such as pheromones, kairmones, aposematism, mimicry, crypsis or more specialized pollinator plant relationships (Burger et al., 2021). In addition, more studies will be needed to understand how the hyperbolic space is represted in in olfactory processing in the CNS.

## Acknowledgements

We thank Dr. Jürgen Liebig for use of his lab equipment, Dr. Kavita Sharma for her advice regarding EAG recordings, Dr. Chris Jernigan for help with statistical analysis, and Dr. Margaret Henderson for help with preliminary statistical analyses of odorant mixtures. We thank Kevin L. Haight for valuable comments on the first draft of the manuscript. This research was supported by by the Next Generation Networks for Neuroscience Program (Award #2014217), the Aileen Andrew Foundation, the National Science Foundation (NSF) award numbers IIS-1254123 and IOS-1556388 to TOS and 1556337 to BHS. Both awards from IOS resulted from an NSF Ideas Lab ‘Cracking the Olfactory Code’. FPS and ACK were funded by the European Union’s Seventh Framework Program ([FP7/2007-2013] [FP7/2007-2011]) under grant agreement no. 281093.

---

**Supplemental Figure 1:**
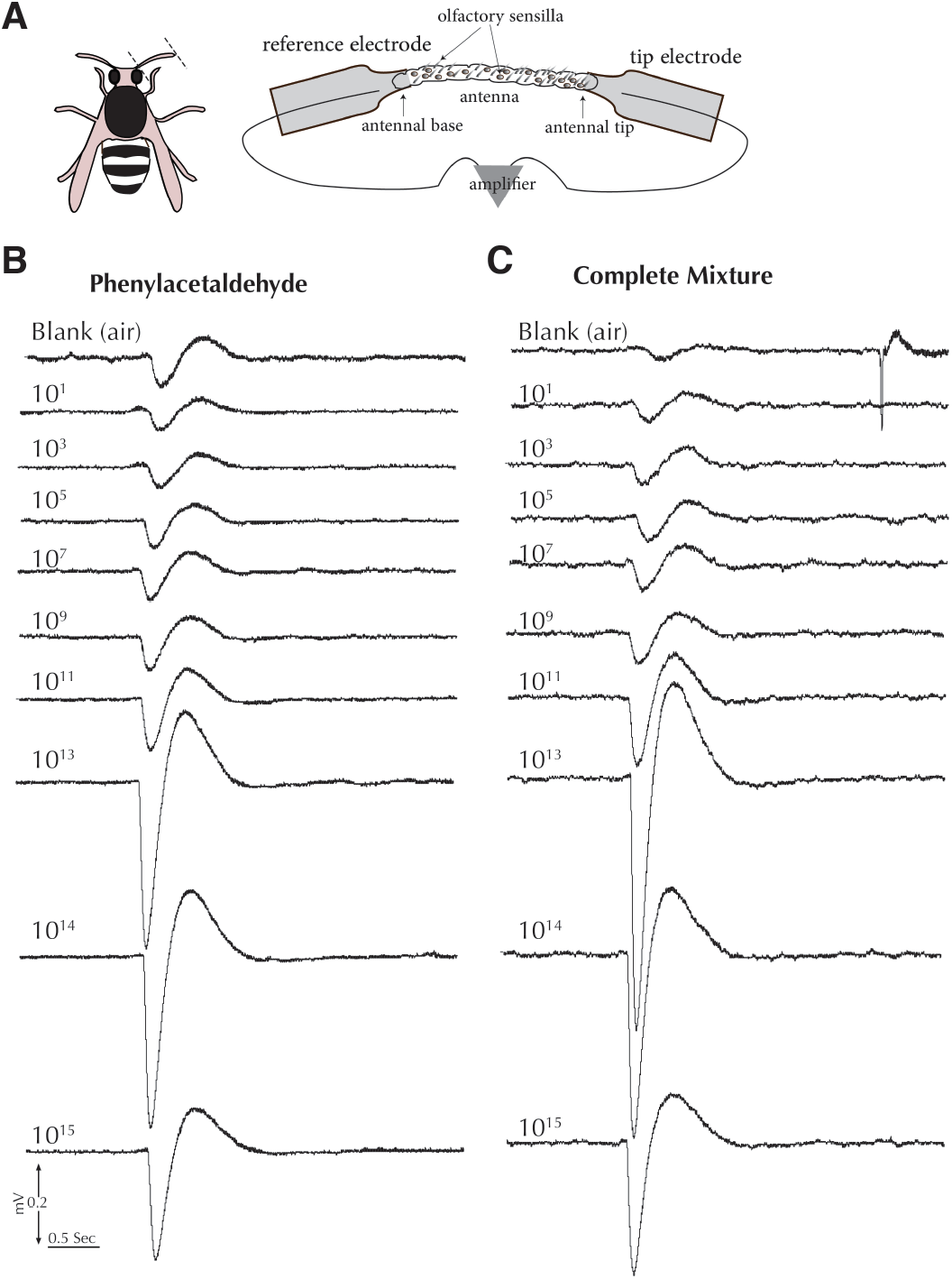
(A) a schematic drawing of an electroantennaography setup. The two ends of the antennal shaft, after being cut off from a bee’s head capsule, were inserted in the two capillary glasses (reference and tip electrodes) containing insect ringer’s solution. For more details about the procedure see Method Details. Sample electroantennogram traces when a honey bee was exposed to the increasing concentrations of (B) phenylacetaldehyde and (C) complete mixture. Note that the olfactory responses to both stimuli at 10^2^ and 10^3^ g/L are lower than those at 10 g/L. The responses are restored after enough inter-delivery time was given to the OSNs (see Figure 1).

**Supplemental Figure 2:**
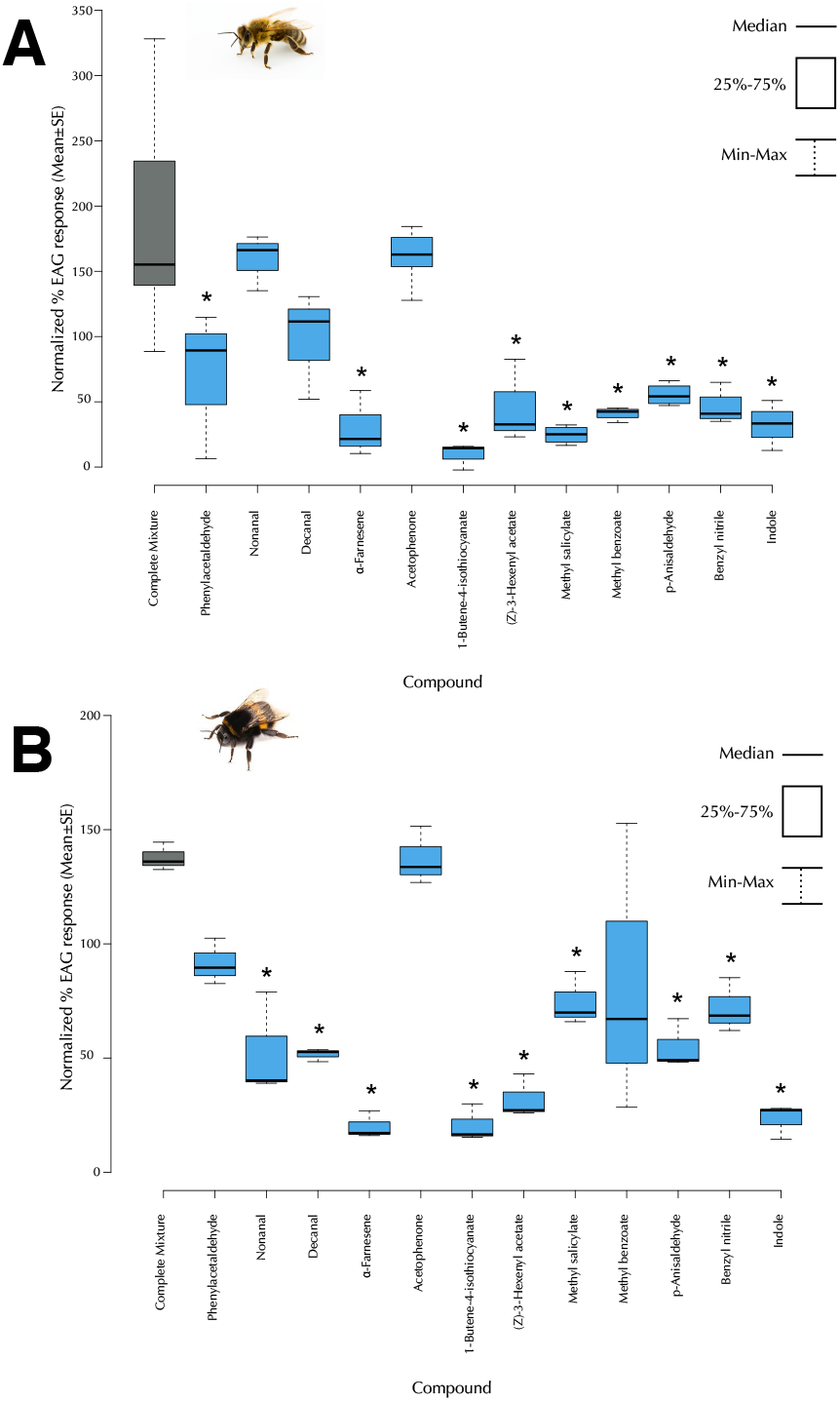
Antennal responses to the individual components and the mixture are different at 10 g/L. Electroantennographic responses of (A) honey bees (n=3) and (B) bumble bees (n=3) to the scent of *Brassica rapa* at 10 g/L. We first compared the intensity of the responses obtained when the compounds were presented individually at 10 g/L and found that the EAG values for the different odors in both species are significantly different (ANOVA, F*_Apis_*= 9.86, F*_Bombus_*= 11.16, df=12, p<0.00001). We then compared the intensity of the responses obtained when the compounds were presented individually at 10 g/L with that in the complete mixture. The EAG values for the different odors in both species are significantly different from those for the mixture (ANOVA, df=12, Tukey’s HSD, p<0.05). Nonetheless, in *Apis mellifera* the response intensity to nonanal, decanal, and acetophenone alone increased to that elicited by the mixture (Tukey’s HSD, p>0.1). In *Bombus terristris*, however, the response to phenylacetaldehyde, acetophenone, and methyl benzoate was similar to the mixture (Tukey’s HSD, p>0.1).

**Supplemental Figure 3:**
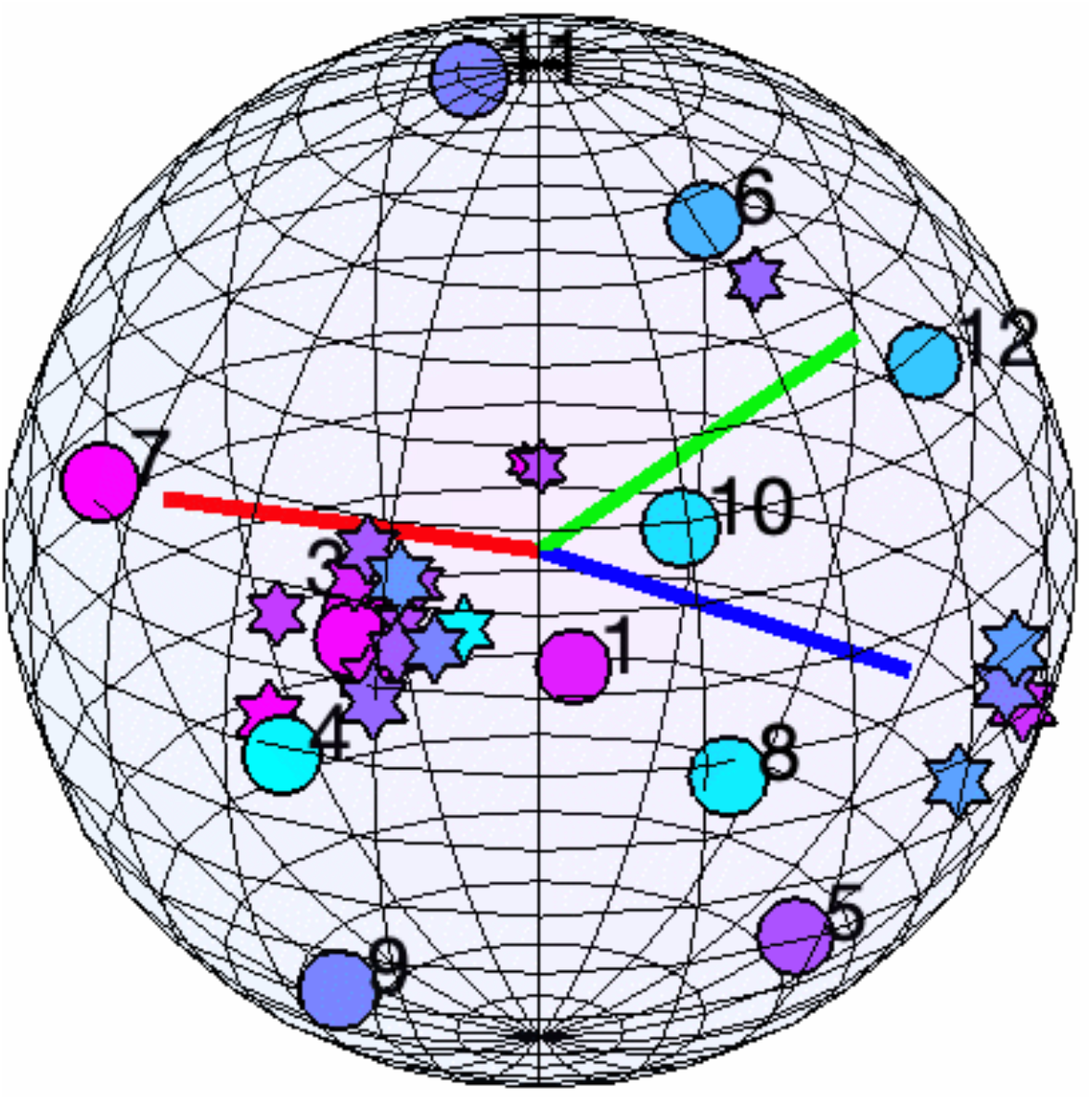
Three dimensional representation of Poincare ball in Fig. 4A.

