## Supplementary figures and images for "Hyperbolic odorant mixtures as a basis for more efficient signaling between flowering plants and bees"

### Fig_Brassica_HMDS_new.gif

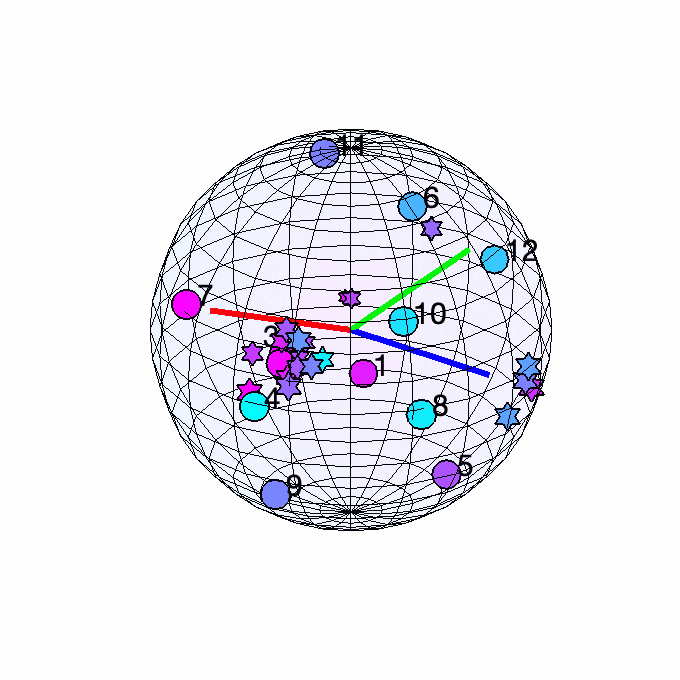
